# Nonlinear locomotion patterns in the Egyptian Locust (Anacridium aegyptium) during walking: a detailed case study

**DOI:** 10.1101/2025.01.18.633722

**Authors:** Arturo Tozzi

**Author notes:** (corresponding author) 1155 Union Circle, #311427 Denton, TX 76203-5017 USA.

## Abstract

We explored the nonlinear movement patterns of *Anacridium aegyptium* during terrestrial locomotion, providing insights into the walking dynamics of this large grasshopper species. Using video recordings, we analysed the trajectory of an insect and quantified key metrics, including curvature, tortuosity and fractal dimension. Curvature analysis revealed irregular turning behaviors with sharp directional changes, suggesting that locomotion was not random but deliberate. Compared with simulated linear trajectories, the curvature exhibited distinct peaks, highlighting the presence of statistically significant nonlinear features in the movement patterns. Phase space reconstruction revealed repetitive patterns indicating the potential presence of a limit cycle attractor. The trajectory remained confined within a specific region of the phase space, highlighting structured dynamics rather than unbounded behaviour. Fractal dimension analysis and Lyapunov exponent were consistent with a stable and predictable system over time, rather than one governed by chaos. These findings align with the behavioral ecology of *A. aegyptium*, suggesting that its walking dynamics are governed by efficient spatial exploration and obstacle negotiation rather than erratic or chaotic motion. Our study underscores the value of advanced mathematical and computational methods in boosting behavioural studies of locomotion. The insights derived from our analysis enhance our understanding of insect locomotion strategies and hold potential applications in the field of biomimetic robotics, where adaptive and efficient movement is mandatory. Future research could explore the impact of environmental factors, such as substrate type and food availability, on the observed nonlinear patterns, providing deeper context to the intricate locomotion behaviour of *Anacridium aegyptium*.

## INTRODUCTION

Locomotion is a fundamental aspect of animal behavior serving crucial roles in habitat navigation, foraging and predator avoidance. The six-legged insects display a remarkable diversity of locomotion strategies, which have been a focal point of neurobiological research for over a century (Bidaye et al, 2018; Heckenthaler et al., 2023; Regeler et al., 2023). Insect locomotion has been extensively studied in the contexts of flying, walking, crawling and central pattern generation, both in natural systems and artificial walking systems (Seipel et al., 2004; Imirzian et al., 2019; Mantziaris et al., 2020; Sabattini et al., 2023). Research on insect locomotion has predominantly concentrated on the biomechanics of jumping and the dynamics of flight. Indeed, recent advancements have introduced a diverse array of modelling and simulation techniques, alongside experimental setups, to investigate collective insect motion. These approaches range from discrete agent-based models of self-propelled particles to continuous frameworks using integral-differential equations (Ariel and Ayali, 2015; Bleichman et al., 2024; Aidan et al. 2024). Grasshoppers are particularly noteworthy for their efficient jumping and walking capabilities. Among these, *Anacridium aegyptium*, commonly known as the Egyptian Locust, stands out due to its large size, robust anatomy and widespread distribution across Mediterranean and subtropical regions. The act of jumping in grasshoppers has garnered significant attention from researchers due to its intricate biomechanics and critical role in their survival strategies (Hawlena et al., 2010: Hawkes et al., 2022). Conversely, solitary walking behaviors in grasshoppers remain relatively underexplored, particularly with respect to nonlinear patterns and their ecological significance.

Nonlinear locomotion encompasses movement patterns deviating from straightforward linear trajectories, often distinguished by irregular pathways (Campos et al., 2010; Xu et al., 2023). Zigzagging paths may optimize resource exploration, while abrupt directional changes might indicate evasive maneuvers against predators. Nonlinear time-periodic models of flight dynamics have been studied, for instance, in the desert locusts *Schistocerca gregaria* (Taylor et al., 2005). Investigating these patterns in *A. aegyptium* could provide valuable insights into the broader principles governing insect locomotion and their potential applications in fields such as robotics and ecological modeling.

Advanced mathematical tools have transformed the study of animal movement, enabling precise quantification of complex trajectories. For instance, custom tracking algorithms have been employed to uncover fundamental animal-animal interactions that drive collective motion in swarms of marching locust nymphs (Ariel et al., 2014). Metrics such as curvature, fractal dimension, tortuosity and Lyapunov exponents are now routinely employed to uncover previously imperceptible locomotion patterns in both natural and artificial systems (Kearns et al., 2027; Suryanto et al., 2022; Xu et al., 2023). Curvature analysis highlights turning behaviors and their frequency, while fractal dimension quantifies the geometric complexity of a trajectory. The Lyapunov exponent, a measure of the sensitivity of a system to initial conditions, can indicate whether movements exhibit chaotic properties (Mehdizadeh, 2018).

This study aims to investigate the nonlinear walking dynamics of *A. aegyptium* using video analysis and advanced computational methods. By quantifying the above-mentioned nonlinear metrics, we seek to characterize the insect’s locomotion patterns and assess their ecological significance. Specifically, we hypothesize that *A. aegyptium* exhibits significant nonlinear features in its walking trajectory, indicative of adaptive and intentional locomotion strategies. In the following sections, we detail the materials and methods used to collect and analyze data, present the results of our quantitative analyses and discuss their implications in the context of both insect ecology and applied sciences. Through this investigation, we aim to bridge the gap between descriptive studies of insect movement and the rigorous mathematical frameworks needed to understand its underlying dynamics.

## MATERIALS AND METHODS

To investigate the nonlinear walking dynamics of *Anacridium aegyptium*, a combination of field observations, video recording and computational analysis was employed. The study focused on an adult male specimen of *A. aegyptium*, casually encountered in its natural environment under undisturbed conditions. The specimen was observed in the Mediterranean region while walking on a glass window. The glass window provided a clear substrate for tracking the insect’s movements, while natural daylight ensured optimal visibility without introducing artificial stressors. The study focused on a single individual, providing a brief but valuable snapshot of its behaviour. Since the observation involved a single individual, there was no risk of interference from conspecifics.

*A. aegyptium*, commonly known as the Egyptian Grasshopper or Egyptian Locust, is a large species in the Acrididae family, commonly found in the Mediterranean basin. It is one of the largest grasshoppers in Europe. It is primarily found perched on trees and shrubs, relying on short flights and jumping for movement. Key identifiers in our study included the distinctive vertical stripes on the eyes, the hind tibiae adorned with two rows of white spines tipped in black and the presence of eight abdominal segments in males, as opposed to the seven segments typically found in females (Girardie and Granier, 1974).

### Video processing

Video recordings were captured at 30 frames per second using a camera mounted 1.5 meters above the arena. The field of view covered the entire arena, ensuring that the insect’s trajectory could be tracked continuously. The raw video footage was processed using a custom Python-based software pipeline. The first step of motion capture involved converting the video into individual frames, which were subsequently analyzed using an object detection algorithm based on Optical flow (Lucas-Kanade method) via OpenCV (Al-Qudah and Yang, 2023). The algorithm identified the position of the center of the body in each frame, generating a sequence of x and y coordinates representing the insect’s trajectory over time (**Figure 1**). This was determined using simple thresholding to identify the largest contour in the image, which typically corresponds to the main body of the insect. The trajectory data were smoothed using a low-pass filter to reduce noise.

**Figure 1.**
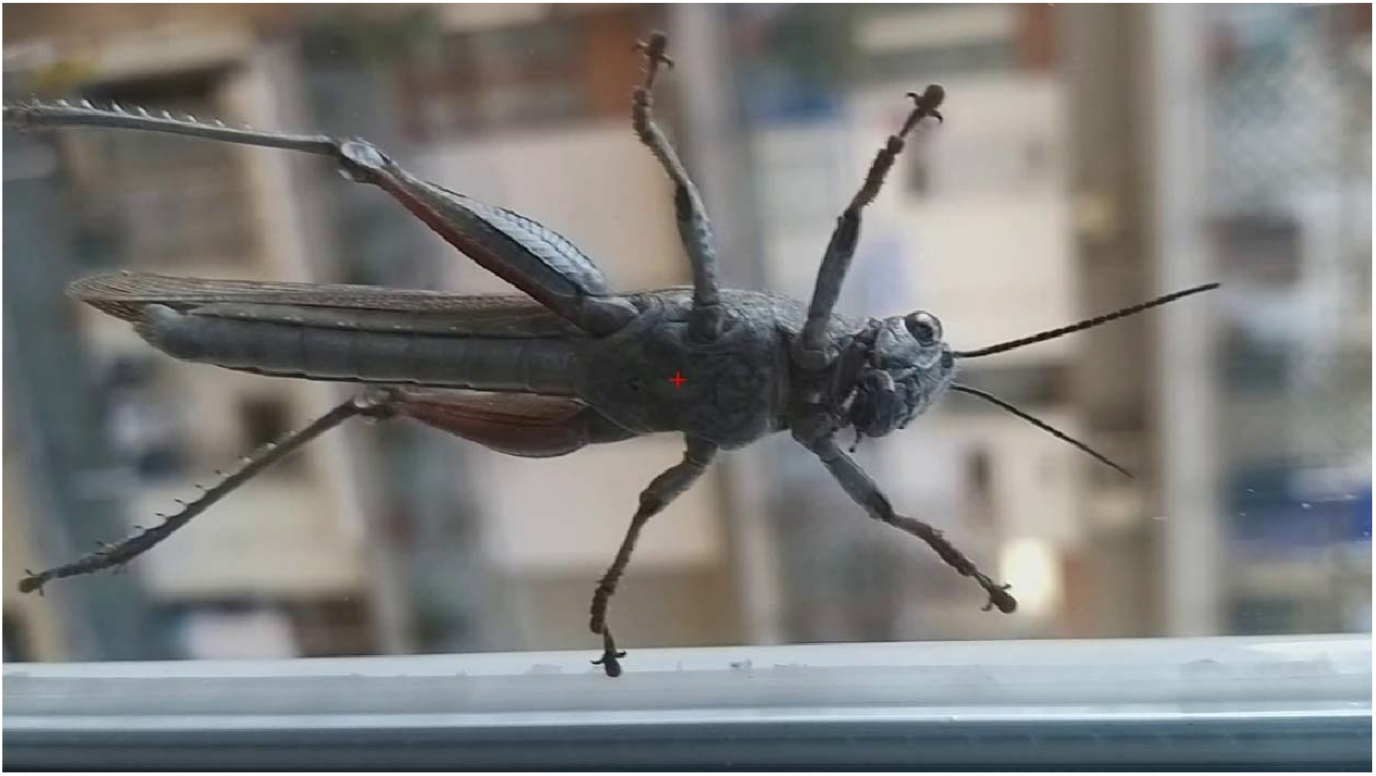
A video frame displaying the detected point on *Anacridium aegyptium*’s body, marked in red. This point was used as the reference for reconstructing the insect’s movement trajectory.

### Nonlinear analysis

Various parameters were derived from the insect’s movement trajectory. **Displacement** represents the distance between the initial and final positions. The **total path length** is the sum of the distances covered between consecutive points along the trajectory. **Average speed** was determined by dividing the total path length by the total time of observation. **Instantaneous speed** was calculated by dividing the distance covered between consecutive points by the time between frames. Finally, the **direction of movement** was measured as the angle of motion between consecutive points, providing insight into the insect’s orientation during its movement.

Next, a range of metrics was calculated to analyse and define the nonlinear movement features. **Curvature** was calculated as the change in direction per unit distance, providing a measure of how sharply the insect turned at each point along its path. **Tortuosity** was assessed as the ratio of the total path length to the straight-line displacement, with higher values indicating more convoluted trajectories. **Fractal dimension**s were estimated using the box-counting method, which involves overlaying a grid of varying box sizes on the trajectory and counting the number of boxes intersected by the path. To further explore the dynamics, **phase space reconstruction** was performed. This involved embedding the trajectory data in a higher-dimensional space using time delays, allowing the identification of patterns that are not apparent in the original two-dimensional trajectory. The embedding dimension and delay time were determined using the false nearest neighbors method and mutual information analysis, respectively (Albers and Hripcsak, 2012; Wallot and Mønster, 2018). The reconstructed phase space was then analyzed for **attractor behavior**, with particular attention to whether the trajectories exhibited features characteristic of limit cycles or other nonlinear dynamic phenomena (Broscheid et al., 2018). Additionally, the Lyapunov exponent was calculated to evaluate the trajectory’s sensitivity to initial conditions. This process involved tracking the divergence of nearby points in phase space over time. The divergence was then plotted on a logarithmic scale and the exponent was estimated from the slope of the resulting curve. A positive Lyapunov exponent indicated the presence of chaotic behaviour.

To ensure the robustness of the findings, the analyses were repeated using a 5-second subset of the data, applying varying smoothing parameters and detection thresholds. Sensitivity analyses were also conducted to ensure that the results were not unduly influenced by the choice of parameters such as the box size in the fractal dimension analysis or the embedding dimension in the phase space reconstruction.

### Tools and statistical analysis

Computational analyses were implemented in Python using libraries such as NumPy, SciPy and Matplotlib for numerical computation and visualization. Statistical analyses were conducted to test the significance of the observed nonlinear features. Curvature and tortuosity metrics were compared against null models generated from simulated linear trajectories created by randomly sampling points within the arena and interpolating straight-line paths between them. The distributions of curvature and tortuosity in the observed and simulated datasets were compared using Kolmogorov-Smirnov tests. The fractal dimension of the observed trajectories was compared against random walk models to determine whether the observed complexity exceeded that expected from stochastic motion.

## RESULTS

The video analysis of *Anacridium aegyptium*’s walking dynamics, captured at a resolution of 1080 × 1920 pixels (width × height), provided a detailed quantification of the movements, uncovering significant nonlinear features. Over a trajectory spanning 433 frames at approximately 30 frames per second, the insect demonstrated a total path length of approximately 1189 pixels, far surpassing its net displacement of 24 pixels. This discrepancy underscores the pronounced nonlinearity of the insect’s motion, further supported by a tortuosity value of 49.43, which reflects the highly convoluted nature of its path. Comparisons between the total path length and the straight-line displacement further emphasized this nonlinearity, with substantial differences observed between the two measures throughout the trajectory (**Figure 2**).

**Figure 2.**
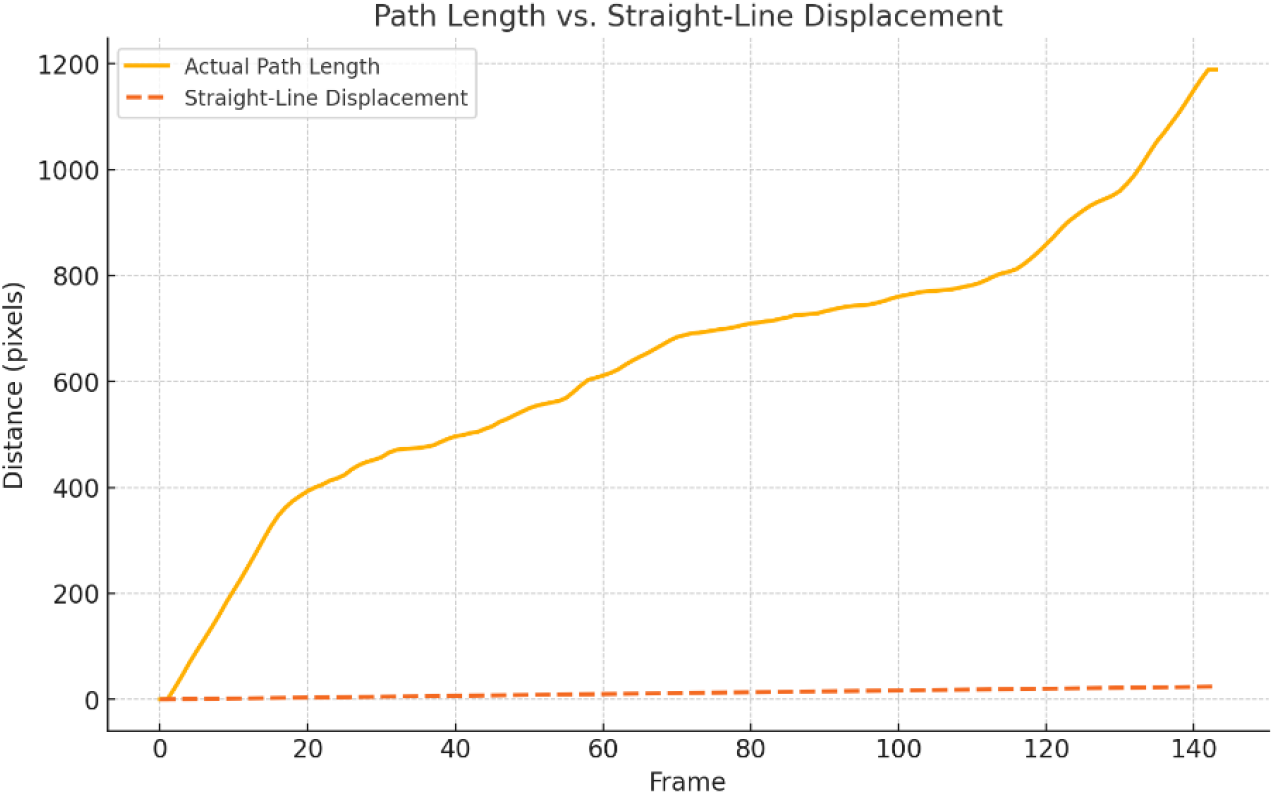
Relationship between the path length and the straight-line displacement of the insect’s trajectory. The solid line represents the cumulative actual path length, capturing the total distance travelled during locomotion. The dashed line reflects the straight-line displacement, indicating the direct distance from the starting point to the endpoint of the trajectory. The notable gap between these two lines serves as clear evidence of the high tortuosity and the nonlinear characteristics inherent in the insect’s movement pattern.

The curvature analysis revealed subtle yet consistent turning behaviors, with sharp directional changes frequently interrupting the trajectory. These patterns suggest that the movement is not random but deliberate, possibly driven by environmental stimuli or internal decision-making processes. Compared with simulated linear trajectories, the curvature exhibited higher variability and distinct peaks, highlighting the presence of statistically significant nonlinear features in the movement patterns (**Figure 3**). This finding confirmed the existence of nonrandom turning behaviors.

**Figure 3.**
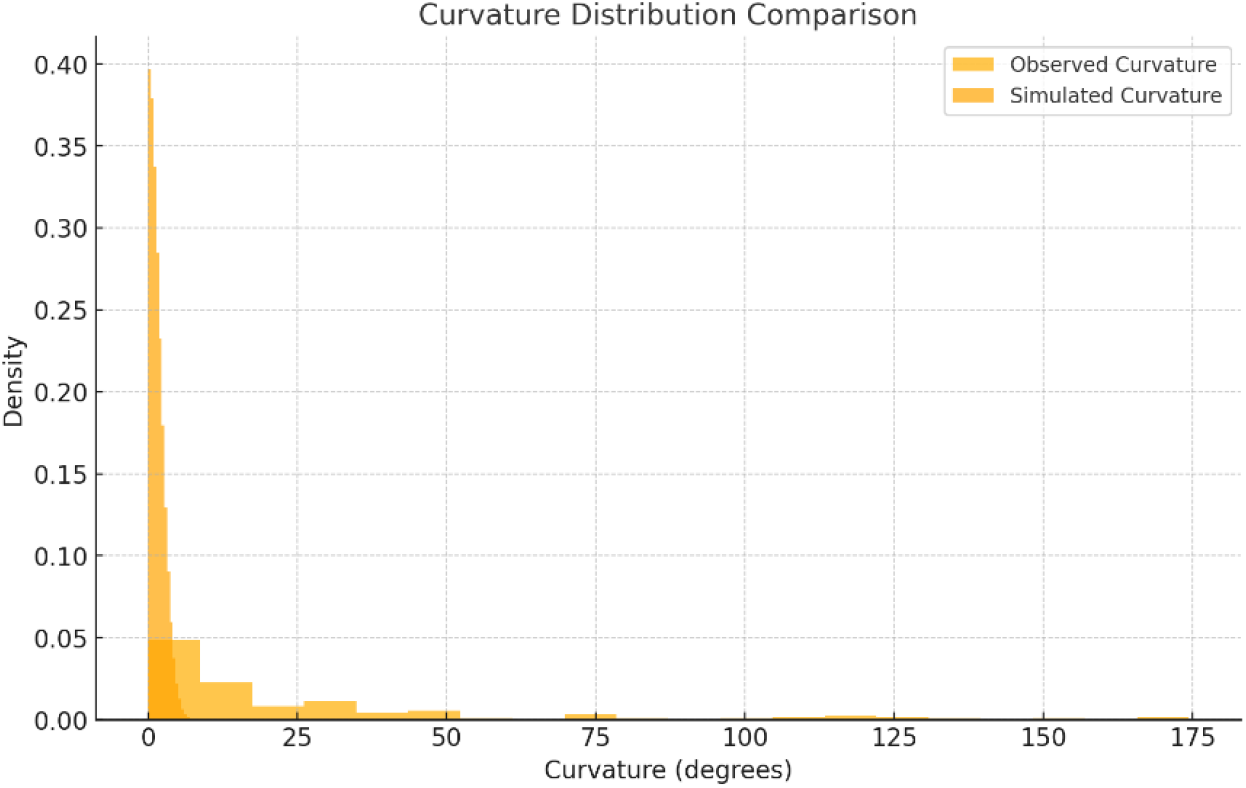
Comparison of curvature distributions. Yellow bars represent the observed curvature values derived from the insect’s trajectory, while orange bars depict the curvature distribution from random simulated trajectories. The observed curvature distribution reveals a statistically significant deviation from the simulated random trajectories.

The analysis of the direction of movement identified dominant frequencies, with direction oscillations occurring at approximately 0.42 Hz (**Figure 4**). This suggested a consistent rhythmic pattern in the insect’s behaviour. In contrast, speed oscillations did not exhibit statistically significant periodicity, indicating that variations in speed may be more context-dependent or driven by external stimuli rather than inherent rhythmicity.

**Figure 4.**
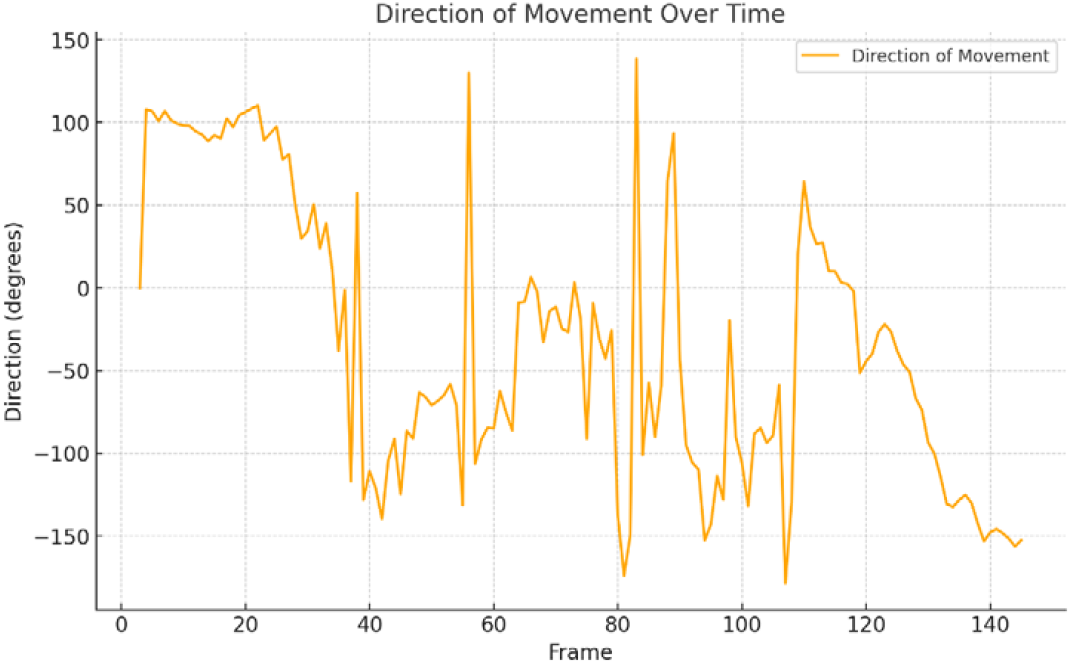
Direction of movement over time. Peaks in the plot correspond to bursts of movement, while dips indicate slower movement or pauses.

**Figure 5.**
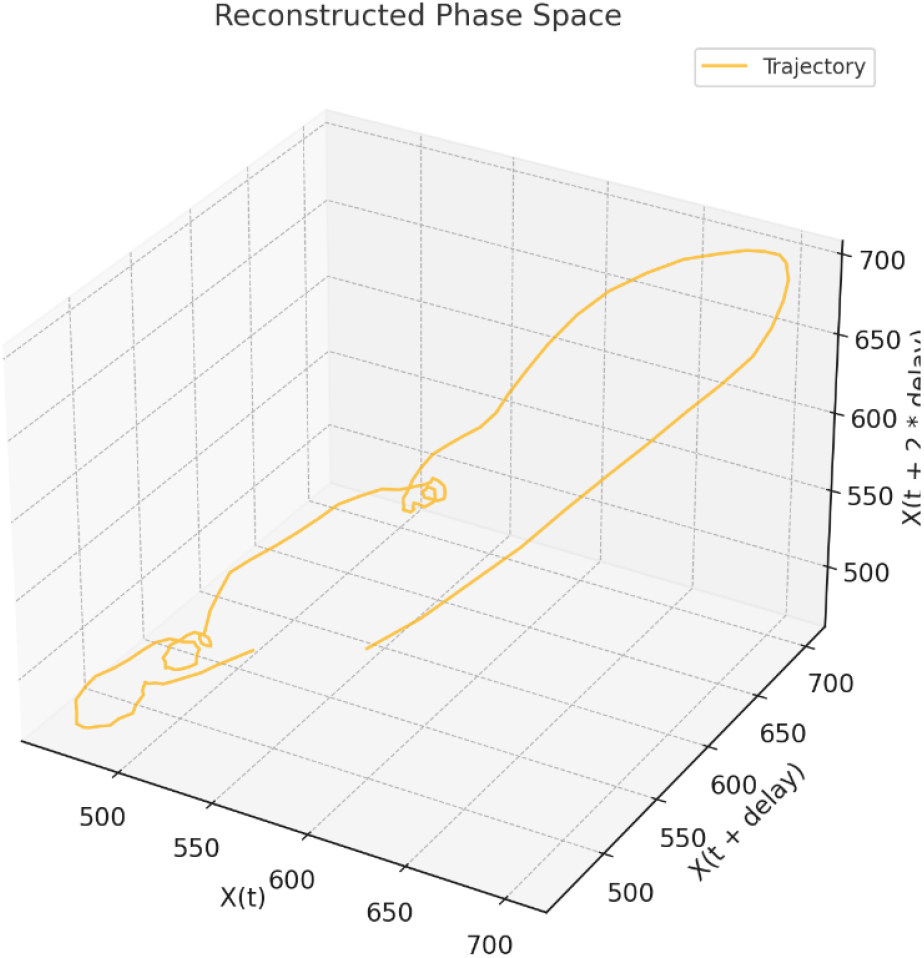
Phase space reconstruction of the 3D trajectory using time-delay embedding. The trajectory does not converge toward a single point, indicating the absence of a fixed-point attractor. Indications of repetitive loops suggest the potential presence of a limit cycle attractor, although the loops lack perfect regularity. The absence of irregular divergence in the phase space suggests that the system does not exhibit chaotic features or a strange attractor.

Phase space reconstruction illustrated the bounded nature of the motion, with repetitive patterns pointing toward a potential limit cycle attractor (**Figure 5**). This attractor behavior indicated a structured yet flexible movement strategy, allowing for environmental adaptability. The trajectory was confined within a bounded region of the phase space, suggesting structured dynamics rather than random or unbounded behaviour. The reconstructed phase space showed clear loops without chaotic divergence, consistent with a stable and predictable system rather than one governed by chaos. The calculated Lyapunov exponent, approximately -7.5 × 10□ □, further supported the absence of chaos, pointing towards convergence rather than divergence in the system’s dynamics. Fractal dimension analysis provided additional insights into the complexity of the movement. With an estimated fractal dimension of 0.73, the trajectory displayed a constrained complexity characteristic of structured but not chaotic motion. This metric underscored the insect’s ability to navigate effectively within defined spatial limits while maintaining a balance between exploration and efficiency.

**Figure 5.**
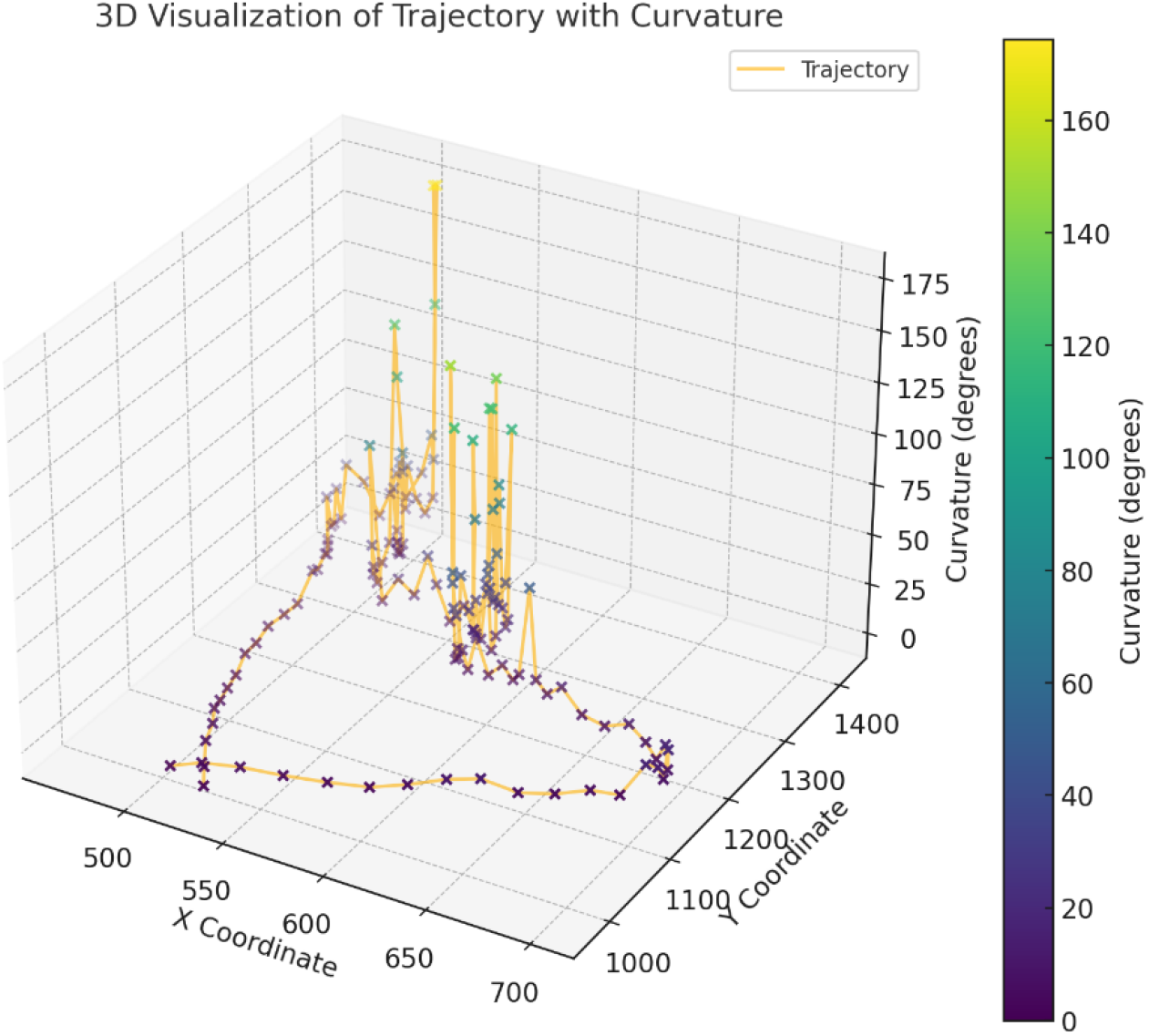
3D visualization of the insect’s trajectory, incorporating curvature as the third dimension. Each point represents the detected position of the tracked feature across frames. The x and y axes represent the spatial movement of the insect while the z-axis corresponds to curvature, with higher values signifying sharper turns. The colour intensity of the points reflects the curvature at each location, with more intense colours highlighting areas where sharper turns occur. This visualization emphasizes the nonlinearity of the trajectory, highlighting deviations and directional changes over time.

Visual analyses complemented these findings. Trajectory plots highlighted regions of high curvature, represented by intensified color gradients, corresponding to areas of sharper turns or more frequent directional adjustments (**Figure 6**). The curvature was nonuniformly distributed, with distinct peaks suggesting localized exploratory behavior or evasive maneuvers.

## CONCLUSIONS

We provide a comprehensive analysis of the nonlinear walking dynamics of *Anacridium aegyptium*. The computational approach employed in this study enabled the extraction of detailed and quantitative insights from behavioral data. The integration of tools such as curvature analysis, fractal dimension estimation and phase space reconstruction allowed for a multidimensional understanding of grasshopper’s locomotion that extends beyond traditional observational methods. Our findings underscore the importance of nonlinear analysis in understanding animal movement, particularly for behaviors that cannot be adequately described by linear models. Nonlinear movement patterns, such as Lévy flights or Brownian motion, are well-documented in insects during behaviors like foraging, mate searching or escape responses (Taylor et al., 2005). However, detailed statistical analysis of curvature remains relatively uncommon. Notably, one of the most striking outcomes of our research is the identification of curvature and tortuosity as key indicators of the insect’s adaptive nonlinear walking behaviour. The significant deviations from random linear trajectories suggest that *A. aegyptium* employs deliberate turning and path-convoluting strategies. These movements may reflect purposeful behaviors, including optimizing foraging efficiency, avoiding obstacles avoidance, searching or responding to potential threats. High curvature could indicate localized exploration near stimuli, while sudden changes may signify evasive actions. Additionally, curvature may reveal biomechanical limitations or energy-efficient turning mechanisms inherent to the insect’s physiology, as frequent turning may incur energy costs that influence resource optimization strategies. Furthermore, significant curvature deviations can serve as reliable indicators for distinguishing insect behaviors from random noise or external disturbances. Novelty can also stem from linking curvature to specific environmental or behavioral contexts, such as food searching or predator avoidance. In the case of *A. aegyptium*, nonlinear movement patterns during walking are of particular interest due to the ecological contexts in which they occur. As a large, terrestrial insect, *A. aegyptium* often traverses uneven substrates, navigates dense vegetation and interacts with potential threats or resources. The frequent directional changes, bounded trajectories and repetitive loops likely represent an optimized trade-off between environmental exploration and energy conservation.

The use of fractal dimension analysis further revealed the constrained complexity of the trajectories, highlighting the ability to navigate within defined spatial boundaries while maintaining efficient movement patterns. Phase space reconstruction and Lyapunov exponent calculations provided additional depth to our understanding of walking dynamics. The identification of a possible limit cycle attractor indicates repetitive and bounded behaviors, which may reflect innate or environmentally influenced patterns. While no evidence of chaotic dynamics was found, the bounded and structured trajectories point to a deterministic system governed by environmental feedback and internal rules. This structured behavior aligns with the ecological requirements of *A. aegyptium*, enabling it to adapt to complex terrains while conserving energy. The bounded movement and lack of chaotic divergence are consistent with efficient navigation strategies in natural habitats. These findings suggest potential applications in biomimetic robotics, where adaptive and efficient movement strategies are critical.

Comparing observed curvature to random trajectories represents a methodological advancement that has been already used in standard ecological studies. Artificially generated video sequences have been introduced, combining known real-animal postures with randomized body positions, orientations and sizes (Arent et al., 2021). The novel contribution of our study is that is rigorously tests curvature against null models such as random linear paths incorporating quantitative metrics like curvature distribution or tortuosity. The novelty is enhanced by the focus on a specific insect species or behavior (i.e., a grasshopper’s movement during natural conditions) that has been scarcely analyzed in this way. The methodologies developed here could be adapted to study other forms of animal movement, from terrestrial vertebrates to aquatic species, providing a versatile toolkit for ecological and biomechanical research. For navigation studies, understanding nonlinear movements may illuminate how insects process sensory information and make decisions. Additionally, recognizing curvature patterns in movement may aid pest control strategies or contribute to ecological monitoring efforts. The insights gained from understanding the nonlinear dynamics of *A. aegyptium*’s walking behavior could inform the design of biomimetic robots capable of adaptive and efficient locomotion in complex terrains (Gart et al., 2018). By mimicking the turning strategies, bounded trajectories and constrained features observed in this study, robotic systems could achieve enhanced agility and robustness.

Despite our findings, several limitations must be acknowledged. First, the experimental setup cannot fully capture the complexities of the insect’s native habitat. Factors present in the wild, such as predation risk, interspecies interactions, and varying substrate types, were not considered in this study. Second, the analysis relied on two-dimensional trajectory data, which may overlook vertical components of movement. Future studies incorporating three-dimensional tracking techniques could provide a more holistic perspective. Whereas marker-based motion capture systems are very robust and easily adjusted to suit different setups, tracked species or body parts, they cannot be applied in experimental situations where markers interfere with natural behavior, e.g., when tracking delicate, elastic or sensitive body structures (Arent et al., 2021). Another limitation is the limited generalizability of the findings, as the observations were based on a single individual. Expanding the sample size and including a broader range of conditions would strengthen the robustness of the conclusions. Additionally, while advanced metrics such as fractal dimension and Lyapunov exponents were calculated, these analyses are sensitive to parameter choices such as time delays and embedding dimensions. Further refinement of these methods could enhance the reliability of future studies. While this research focused primarily on movement patterns, future studies could incorporate additional variables such as environmental. For example, controlled manipulations of environmental features could help disentangle the roles of intrinsic and extrinsic factors in driving observed behaviors. Future directions include linking curvature to environmental factors like light or food availability to determine its drivers, conducting comparative studies across species and behaviors and exploring other nonlinear features such as speed oscillations or pauses.

In conclusion, this interdisciplinary investigation into the nonlinear walking dynamics of *Anacridium aegyptium* demonstrates the value of advanced analytical techniques in uncovering hidden patterns and address fundamental questions about movement and behavior. Despite its limitations, this study offers a compelling framework for examining locomotion through the lens of nonlinear dynamics, with applications ranging from ecology and evolutionary biology to biomechanics and robotics.

## DECLARATIONS

### Ethics approval and consent to participate

This research does not contain any studies with human participants or animals performed by the Author.

### Consent for publication

The Author transfers all copyright ownership, in the event the work is published. The undersigned author warrants that the article is original, does not infringe on any copyright or other proprietary right of any third part, is not under consideration by another journal, and has not been previously published.

### Availability of data and materials

all data and materials generated or analyzed during this study are included in the manuscript. The Author had full access to all the data in the study and take responsibility for the integrity of the data and the accuracy of the data analysis.

## Acknowledgements

none.

## Authors’ contributions

The Author performed: study concept and design, acquisition of data, analysis and interpretation of data, drafting of the manuscript, critical revision of the manuscript for important intellectual content, statistical analysis, obtained funding, administrative, technical, and material support, study supervision.

### Declaration of generative AI and AI-assisted technologies in the writing process

During the preparation of this work, the author used ChatGPT to assist with data analysis and manuscript drafting. After using this tool, the author reviewed and edited the content as needed and takes full responsibility for the content of the publication.

